# Mitofusin2 cooperates with Nuage-associated proteins and involves mRNA translational machinery in controlling mRNA fates during spermatogenesis

**DOI:** 10.1101/2020.03.29.015024

**Authors:** Xiaoli Wang, Yujiao Wen, Jin Zhang, Shuangshuang Guo, Congcong Cao, Stephen A Krawetz, Zhibing Zhang, Shuiqiao Yuan

## Abstract

Mitochondria play a critical role in spermatogenesis and regulated by several mitochondrial fusion proteins. Its interaction with other organelles forms several unique structures, including mitochondria-associated ER membrane (MAM) and a specific type of Nuage close to mitochondria. However, the importance of mitochondria functions and mitochondrial fusion proteins in its associated-structure formation and mRNA translation during spermatogenesis remain unclear. Here, we show that Mitofusin 2 (MFN2), a mitochondrial fusion GTPase protein, cooperates with Nuage-associated proteins, including MIWI, DDX4, TDRKH and GASZ and involves translational machinery to control the fates of gamete-specific mRNAs in spermatogenesis. Conditional mutation of *Mfn2* in postnatal germ cells results in male sterility due to germ cell developmental defects characterized by disruption of mitochondrial morphology, abnormal MAMs structure, aberrant mRNA translational processes, and anomalous splicing events. Moreover, MFN2 interacts with MFN1, another mitochondrial fusion protein with high-sequence similar to MFN2, in testes to facilitate spermatogenesis. Mutation of *Mfn1* and *Mfn2* simultaneously in testes causes very severe infertile phenotypes. Importantly, we further show that MFN2 is enriched in polysome fractions in testes and interacts with MSY2, a germ cell-specific DNA/RNA-binding protein, and eukaryotic elongation factor 1 alpha (eEF1A) to control gamete-specific mRNA translational delay during spermatogenesis. Collectively, our findings demonstrate that MFN2 works with Nuage-associated proteins and involves translational secession to regulate gamete-specific mRNA fates. Our data reveal a novel molecular link among Mitofusins, Nuage-associated proteins, and mRNA translational processes in controlling male germ cell development.

## Introduction

Mitochondria are dynamic organelles in which maintained by a balance between fusion and fission in mammals. They play crucial roles in multiple biological processes, including energy generation, calcium homeostasis, signal transduction, and apoptosis(Chen and Chan, 2004; Westermann, 2010). In mammalian testes, the shape of germ cell mitochondria changes markedly during spermatogenesis, of which there are three main types. It displays the orthodox-type in Sertoli cells, spermatogonia, and preleptotene/leptotene spermatocytes. While the intermediate-type in zygotene spermatocytes and the condensed type in pachytene spermatocytes and early spermatids. In late spermatids and spermatozoa, it shifts back to the intermediate-type (De Martino et al., 1979; Ramalho-Santos et al., 2009). Emerging evidence indicates that mitochondrial function is a critical determinant of male germ cell development(Zhang et al., 2016; Ren et al., 2019).

Additionally, the mitochondria and endoplasmic reticulum (ER) form contacts called mitochondria-associated ER membranes (MAMs), which are important for supporting essential structural and functional inter-organelle communication including calcium and phospholipid exchange, mitochondrial biogenesis, autophagy, ER stress and unfolded protein response (UPR)(Paillusson et al., 2016). Alteration of MAMs increasingly was reported in several human diseases, highlighted by neurodegenerative diseases(Paillusson et al., 2016). Our previous work has demonstrated an abundance of MAMs in mouse and human testes and identified a large portion of MAM proteins in testes (Wang et al., 2018), which suggests that MAM proteins may have critical functions in spermatogenesis.

Mitofusins (MFN1/2), the homologs of Fzo in yeast and Drosophila, are the critical regulators of mitochondrial fusion in mammalian cells and enriched in MAMs (Hales and Fuller, 1997; Santel and Fuller, 2001; Wang et al., 2018). Loss of either *Mfn1 or Mfn2* in mice leads to embryonic lethality, and cultured cells obtained from the mutant mouse embryos display overtly fragmented mitochondria (Chen et al., 2003). Specific ablation of *Mfn2* in anorexigenic pro-opiomelanocortin (POMC) neurons in the hypothalamus results in aberrant MAMs, defective POMC processing, ER stress-induced leptin resistance, hyperphagia, reduced energy expenditure, and obesity (Schneeberger et al., 2013). Interestingly, the mutations in human MFN2, but not MFN1, lead to Charcot–Marie–Tooth disease type 2A, a neurodegenerative disorder characterized by progressive sensory and motor losses in the limbs (Zuchner et al., 2004; Chen et al., 2007; Misko et al., 2012; Rouzier et al., 2012; Bouhy and Timmerman, 2013). Additionally, conditional knockout *Mfn1,* but not *Mfn2,* in growing oocytes results in female infertility(Hou et al., 2019). Although MFN1/2 deficiency in male germ cells leads to mitochondrial defects and male infertility(Zhang et al., 2016; Varuzhanyan et al., 2019), the underlying mechanism of how Mitofusins regulate spermatogenesis and male germ cell development remains largely unknown.

During spermatogenesis, multiple specific mitochondria-associated germinal structures are present in the cytoplasm of male germ cells, termed Nuage, which are electron-dense, non-membranous, close to mitochondria, variant size yielding to different protein components, including inter-mitochondrial cement (IMC), piP-body, and chromatoid body (CB)(Wang et al., 2020). Several proteins have identified to localize the Nuage, such as MIWI, DDX4, MAEL, TDRKH, and GASZ(Wang et al., 2020). Interestingly, MIWI, DDX4, and MAEL are expressed in both IMC and CB, while TDRKH and GASZ localize only to the IMC (Toyooka et al., 2000; Ma et al., 2009; Saxe et al., 2013; Takebe et al., 2013). The Nuage is the proposed sites of germ cell functions with multiple RNA processing events, including translational regulation, RNA-mediated gene silencing, mRNA degradation, piRNA biogenesis, and nonsense-mediated mRNA decay(Kotaja and Sassone-Corsi, 2007; Wang et al., 2020). Besides, spermatogenesis undergoes two rounds of transcriptional secessions: one is during the meiosis, which includes the synapsis and desynapsis, and the other is the histone-protamine transition process during late spermiogenesis(Sassone-Corsi, 2002). During the transcriptional secessions, a large number of germ cell-specific mRNAs exhibit translational repression with a lag of up to a week between their transcription and translation(Iguchi et al., 2006). The evidence for Nuage regulation of mRNA processing is strongest for the chromatoid body. Translational regulatory proteins such as MIWI, TDRD6, MAEL have been localized to the chromatoid body and associated with mRNA translational machinery and piRNA biogenesis(Grivna et al., 2006; Vasileva et al., 2009; Takebe et al., 2013; Castaneda et al., 2014; Fanourgakis et al., 2016). Thus, multiple components necessary for transcript processing physically connected to the Nuage during spermatogenesis. Transcriptional and post-transcriptional regulation in testis is mainly due to activation/inactivation of specific gene expression programs in meiotic spermatocytes and post-meiotic round spermatids. However, knowledge about the specific regulators and underlying mechanisms in such programs during germ cell development is limited.

In this study, we report that MFN2, a mitochondrial fusion GTPase protein, interacts with Nuage-associated proteins, including MIWI, DDX4, GASZ, and TDRKH in the testes and associates with translational machinery to regulate male germ cell development and spermatogenesis. We discovered that MFN2 provides a critical component to the translational machinery while interacting with the Nuage-associated proteins in the male germ cells. Loss of function of MFN2 in postnatal germ cells led to male sterility with defects in mitochondrial formation, MAMs structure maintenance, translational processes, and mRNA splicing events. Moreover, we found that MFN2 interoperates with MFN1, another mitochondrial fusion protein with high sequence similarity to MFN2, in testes to facilitate spermatogenesis. Simultaneous mutation of *Mfn1* and *Mfn2* in testes causes severely disrupted phenotypes. Inspiringly, our data revealed that MFN2 also cooperates with the germ cell-specific DNA/RNA binding protein MSY2 and eukaryotic elongation factor 1 alpha (eEF1A1 and eEF1A2) to regulate the transcription and translation of MSY2-bound gamete mRNAs in male germ cells. Our study, for the first time, presented below unveils a novel role for MFN2 in mitochondria-associated germinal structure formation and mRNA translational machinery during male germ cell development. The findings revealed a unique mechanism by which MFN2 associates with translational machinery through the Nuage-associated proteins, MSY2 and eukaryotic elongation factor 1 alpha (eEF1A1 and eEF1A2) to control male gamete-specific mRNA fates by maintaining its storage/translational delay in the germline.

## Results

### Expression and localization of MFN2 in the male germ cells during spermatogenesis

Although MFN2 is a ubiquitously expressed protein located on both the outer mitochondrial membrane (OMM) and the ER surface in mammalian cells (de Brito and Scorrano, 2008), to date, no information is available on the pattern of its expression in male germ cells during spermatogenesis. By performing RT-qPCR and Western blot assays, we found that both mRNA and protein levels of the *Mfn2* gene preferentially and highly expressed in mouse testes (Supplementary Figure.S1A-B). We then analyzed the expression of *Mfn2* in testes at various postnatal ages. The results showed that both mRNA and protein are expressed from postnatal day 0 (P0) testes to adult testes but strongly accumulate from P14 when the testes mainly consist of spermatocytes (Supplementary Figure.S1C-D). Immunofluorescence staining of MFN2 in developing testes further revealed that MFN2 mostly expressed in spermatogonia at P0 and P7, and finally highly expressed in late pachytene spermatocytes and round spermatids at P14 to P28 (Supplementary Figure.S1E).

We next determined the subcellular localization of MFN2 in adult testes during spermatogenesis by co-staining MFN2 with γ-H2AX (a marker of meiotic DNA damage response). We observed the presence of MFN2 throughout most of the germ cell development, including the mitotic spermatogonia, meiotic spermatocytes (pre-leptotene to diplotene), and round spermatids (Figure.1A). Interestingly, we found that MFN2 expressed in the cytoplasm of spermatogenic cells with the highest expression level in the late-pachytene spermatocytes and round spermatid with a granular cytoplasmic localization. Modest cytoplasmic expression in meiotic metaphase spermatocytes, followed by spermatogonia, preleptotene, leptotene, and zygotene spermatocytes, was also observed (Figure.1A-B). Moreover, both mRNA and protein levels of *Mfn2* were highly expressed in pre-meiotic spermatocytes and post-meiotic round spermatids while lowly expressed in Sertoli cells (Supplementary Figure.S1F-H). Since *Mfn2* encodes a mitochondrial outer membrane protein, we confirmed its co-localization with ATP5A (an outer mitochondria membrane marker) in adult testes by co-immunofluorescence staining (Figure.1C). Together, these data indicate that MFN2 highly expressed in the cytoplasmic granules of pre-meiotic spermatocytes and post-meiotic round spermatids and may play an essential role in spermatogenesis.

**Figure. 1.**
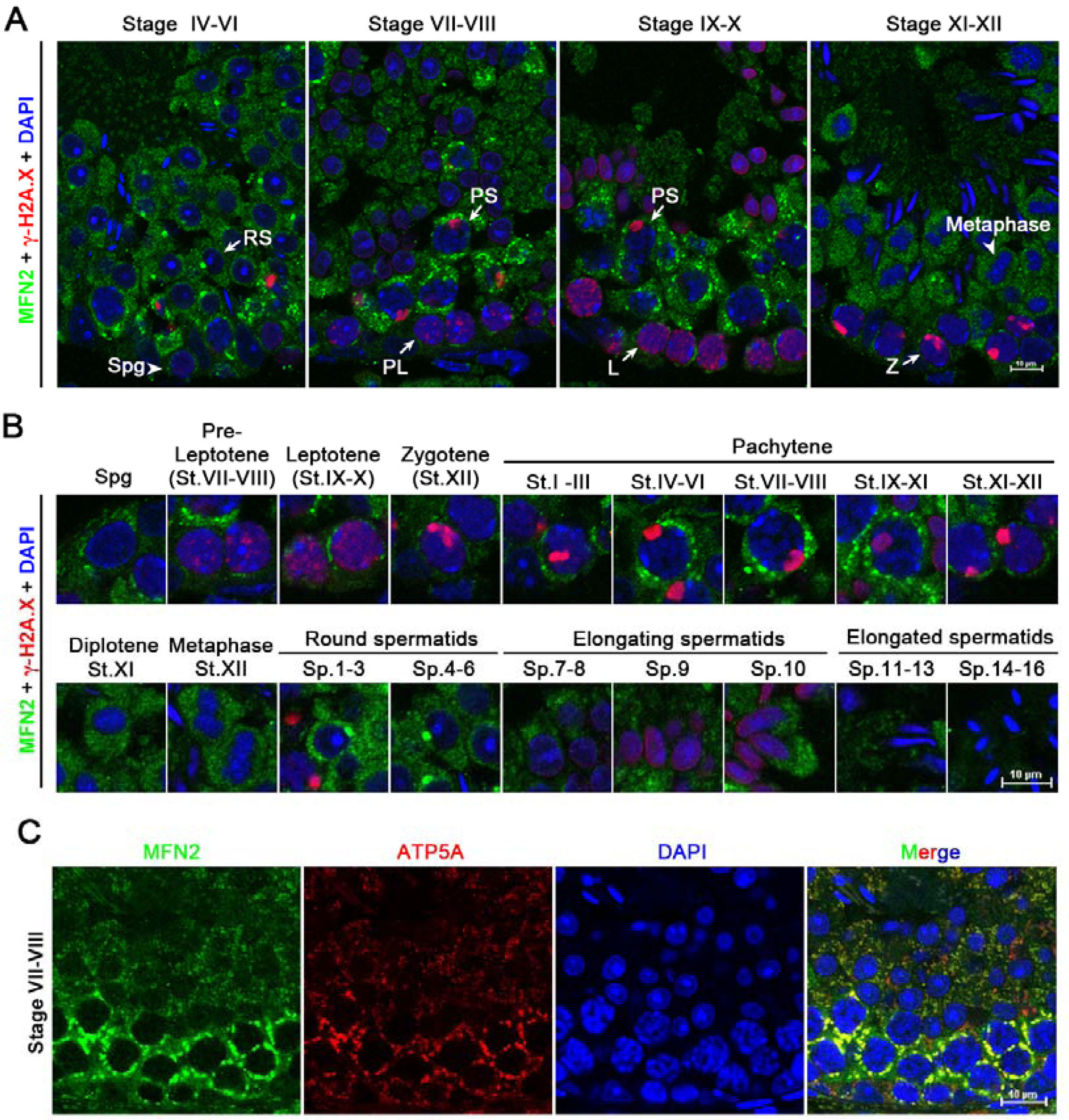
MFN2 displays a dynamic cytoplasm expression profile during spermatogenesis. (**A**) Double immunostaining with MFN2 and γ-H2A.X on WT (wild-type) adult testis sections in different stages of seminiferous tubules are shown. Nuclei were stained with DAPI. Scale bar=10µm. Spg, Spermatogonia; PL, Preleptotene spermatocytes; L, Leptotene spermatocytes; Z, Zygotene spermatocytes; PS, Pachytene spermatocytes; RS, round spermatids. (**B**) Double immunostaining with MFN2 and γ-H2A.X on different types of spermatogenic cells showing a dynamic expression through the whole process of spermatogenesis. Nuclei were stained with DAPI. Scale bar=10µm. (**C**) Co-immunostaining of MFN2 and ATP5A (an outer mitochondria membrane marker) in stage VII-VIII seminiferous tubule from WT testes. Nuclei were stained with DAPI. Scale bar=10µm.

### Loss of MFN2 in postnatal germ cells leads to male sterility

To elucidate the physiological role of MFN2 in spermatogenesis, we generated germline-specific knockout mice by using *Stra8*-Cre transgenic mice in which Cre is expressed in differentiating spermatogonia (Sadate-Ngatchou et al., 2008) to delete exon 6 of *Mfn2* gene (Supplementary Figure.S2A). The efficiency of this *Mfn2* conditional knockout mouse (*Stra8-Cre; Mfn2^loxp/Del^*, herein called *Mfn2*-cKO) was verified by determining *Mfn2* mRNA and protein levels in testes. Both *Mfn2* mRNA and protein levels dramatically reduced in *Mfn2*-cKO testes compared to that of control testes (Figure.2A-D). Thus, we successfully generated the postnatal male germ cell-specific knockout *Mfn2* mutants.

**Figure. 2.**
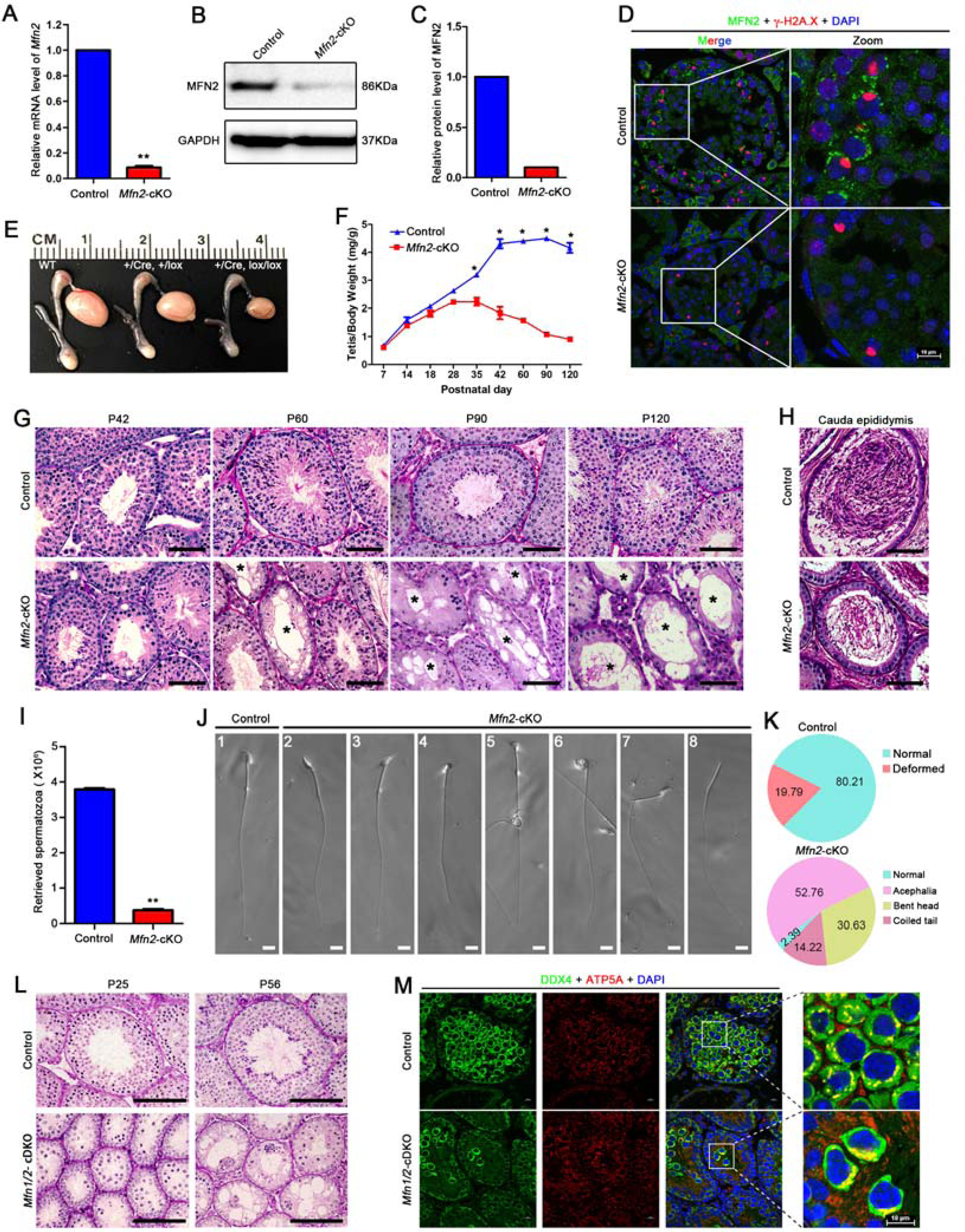
Conditional ablation of MFN2 in postnatal male germ cells results in age-dependent spermatogenic defects and male infertility. (**A**) RT-qPCR analysis showing the dramatic decrease in the mRNA level of *Mfn2* in adult *Mfn2*-cKO testes. Data are presented as mean ± SEM, n = 5. \*\**P*<0.01. (**B**) Western blots of MFN2 expression in control and *Mfn2*-cKO testes. GAPDH serves as a loading control. (**C**) Histogram showing the quantification of MFN2 protein levels in (b). (**D**) Double immunostaining with MFN2 and γ-H2A.X showing MFN2 was undetectable in *Mfn2*-cKO testis sections. Nuclei were stained with DAPI. Scale bar=10µm. (**E**) Gross morphology of the testis and the epididymis from WT, heterozygote (Cre/+, *Mfn2*^+/flox^), and *Mfn2*-cKO (Cre/+, *Mfn2*^flox/flox^) mice at postnatal day 56 (P56). (**F**) Testes growth curve shows that the *Mfn2*-cKO testis weight was significantly decreased from P35. Data are presented as mean ± SEM, n = 5. \**P*<0.05. (**G**) Periodic acid-Schiff (PAS) staining showing the histology of testis sections from control and *Mfn2*-cKO mice at P42, P60, P90 and P120. Asterisk (*) represents the vacuolated seminiferous tubules. Scale bar=50µm. (**H**) PAS staining showing the histology of epididymis sections from control and *Mfn2*-cKO mice at P60. Scale bar=50µm. (**I**) Histograms showing the number of retrieved sperm from the control and *Mfn2*-cKO cauda epididymides. (**J**) Deformed sperm are shown in *Mfn2*-cKO male mice. Scale bar=10µm. (**K**) Pie charts showing the proportional distribution of normal and malformed sperm in control and *Mfn2*-KO male mice. (**L**) PAS staining showing the histology of testis sections from control and *Mfn1/2*-cDKO (*Mfn1* and *Mfn2* double conditional knockout) mice at P25 and P56. Scale bar=100µm. (**M**) Co-immunostaining of DDX4 (a germ cell marker) and ATP5A (a mitochondria marker) in testis sections from control and *Mfn1/2*-cDKO mice at P25. Nuclei were stained with DAPI. Scale bar=10µm.

While *Mfn2*-cKO mice were viable and appeared to be grossly normal, they displayed complete sterility after a 5 month-period fecundity test. Consistent with this infertile phenotype, testis sizes from *Mfn2*-cKO mice were significantly smaller than their control littermates (Figure.2E). The ratio of testis weight/body weight of *Mfn2*-cKO mice was decreased substantially at various ages, starting from P35 to P120 compared with controls (Figure.2F). Histological analyses showed that as the adult *Mfn2*-cKO mice aged, the number of severely atrophic abnormal seminiferous tubules increased (Figure.2G), suggesting that spermatogenic defects are age-dependent in *Mfn2*-cKO mice. Interestingly, when the first wave of spermatogenesis culminated at P42, the late-stage spermatocytes and round spermatids continually decreased in number and appeared to vacuolize gradually (Figure.2G). In comparison, no discernible abnormality found in the *Mfn2*-cKO testes from P7 to P28 (Supplementary Figure.S2B). Consistent with these histological results, the TUNEL assay revealed that the number of apoptotic cells in *Mfn2*-cKO testes at P14 to P28 was comparable with control testes but increased significantly at P35 and P56 (Supplementary Figure.S2C-D). Moreover, the number of spermatozoa retrieved from adult *Mfn2*-cKO cauda epididymis was dramatically reduced compared to that of controls (Figure.2H-I). Only ∼2% of *Mfn2*-cKO epididymal sperm showed normal morphology, compared with ∼80% of the sperm in controls (Figure.2J-K). These data suggest that, upon *Mfn2* deletion in postnatal testes, germ cells gradually lost from P35 by apoptosis, leading to infertility.

### MFN2 interacts with MFN1 in testes and contributes to the male fertility

Since MFN1 and MFN2 are the two Fzo homologs in humans and mice, which interact with each other in mammalian cells coordinately to regulate mitochondrial fusion (Chen et al., 2003), we next asked whether there are interaction and synergistic effects between MFN1 and MFN2 in spermatogenesis. We first examined their interaction effects in adult mouse testes by co-immunoprecipitation (Co-IP). In the MFN2 antibody immunoprecipitants, MFN1 was detected in testes, and in the MFN1 antibody immunoprecipitants, MFN2 was detected as well (Supplementary Figure.S3A). These results confirmed that MFN1 and MFN2 are indeed bona fide interacting partners in the testes. Besides, both MAEL (a piP-body component in the piRNA biogenesis pathway (Soper et al., 2008; Takebe et al., 2013)) and GAPDH were undetectable in MFN2 antibody immunoprecipitants (Supplementary Figure.S3B), confirming the high specificity of MFN2 antibody in Co-IP assays. To determine the roles of MFN1 during spermatogenesis and male germ cell development, we then generated *Stra8*-Cre mediated *Mfn1* conditional knockout mice through the deletion of exon 4 (*Stra8-Cre; Mfn1^loxp/Del^*, herein called *Mfn1*-cKO) in postnatal germ cells (Supplementary Figure.S3C). As expected, *Mfn1*-cKO mice are also infertile due to the disruption of spermatogenesis, which showed smaller testes, decreased testis weight, and disrupted seminiferous tubule structure (Supplementary Figure.S3D-G). However, differing from the age-dependent phenotype of *Mfn2*-cKO mice, *Mfn1*-cKO testes started to exhibit vacuolization defects as early as P28. In comparison, the first wave of spermatogenesis in *Mfn2*-cKO testes proceeds until P42. This disparity between *Mfn1*-cKO and *Mfn2*-cKO mice phenotypes suggests that both MFN2 and MFN1 are essential but play distinct roles for spermatogenesis and male germ cell development.

To better understand the physiological roles of Mitofusins in spermatogenesis and male germ cell development, we next simultaneous deleted *Mfn1* and *Mfn2* in mouse testis using *Stra8*-Cre line mediated Cre-Loxp strategy to create double knockout *Mfn1/2* mice (*Stra8-Cre; Mfn1^loxp/Del^*-*Mfn2^loxp/Del^*, hereinafter referred to as *Mfn1/2*-cDKO). Notably, the testicular disruption in *Mfn1/2*-cDKO mice was much more severe than in either single *Mfn1* or *Mfn2*-cKO mice (Figure. 2L). Compared with the control mice, the diameters of seminiferous tubules from *Mfn1/2*-cDKO mice were much smaller as early as P14 (Supplementary Figure.S4A-B). No round spermatids were observed in *Mfn1/2*-cDKO testes at P25. In comparison, both pachytene spermatocytes and round spermatids were observed in control testes as meiosis completed during first wave spermatogenesis (Figure.2L and Supplementary Figure.S4A). These data suggest that *Mfn1/2*-cDKO mice failed to complete meiosis. Consistent with the phenotypes, testis size of *Mfn1/2*-cDKO adult mice was much smaller than that of *Mfn1* or *Mfn2* single cKO mice (Supplementary Figure.S4C-D). Histological analyses of adult testicular sections further revealed increased atrophic and vacuolated seminiferous tubules in *Mfn1/2*-cDKO testes compared with single *Mfn1* or *Mfn2*-cKO testes (Supplementary Figure.S4E). Strikingly, unlike the distribution of mitochondria in controls, which uniformly distributed in the cytoplasm like a donut, mitochondria were aggregated to one side of the cytoplasm in the *Mfn1/2*-cDKO spermatocytes at P25 (Supplementary Figure.S4F). However, this mitochondria abnormal distribution was not observed in *Mfn1*-cKO or *Mfn2*-cKO mice (Figure.2M and Supplementary Figure.S4F). Together, these results indicate that MFN2 could cooperate with MFN1 to regulate the distribution of mitochondria in the testes, thereby contributing to spermatogenesis and male germ cell development.

### Ablation of MFN2, but not MFN1 in testes disrupt the MAM and ER structures in male germ cells

Since the ablation of both MFN1 and MFN2 in postnatal germ cells caused spermatogenesis defects and male infertility, we next asked why the germ cells fail to full development. Using a transmission electron microscope (TEM) analysis, we observed that mitochondria exhibit swelling and fragmentation in germ cells from adult *Mfn2*-cKO and/or *Mfn1*-cKO mouse testes (Figure.3A-B and Supplementary Figure.S5A). Interestingly, we observed the IMC structure in *Mfn2*-cKO pachytene spermatocytes, but the thickness of IMC appeared to increase (Figure.3A). We then focused on *Mfn2*-cKO round spermatids to further examine the details of mitochondrial defects by measurement of mitochondria length in round spermatids between control and *Mfn2*-cKO mice. The aspect ratio (AR) (AR=major axis/minor axis of mitochondria) is defined to calculate the length of mitochondria. Although the AR of all mitochondria was not significantly decreased in *Mfn2*-cKO round spermatids (Figure.3C), the AR distribution displayed an increased percentage of short mitochondria with AR≤1.5 compared to that of controls (Figure.3D). These data indicate that developmental defects of male germ cells in both *Mfn1*- and *Mfn2*-cKO mice are mainly due to the fragmentation and swelling of mitochondria in germ cells.

**Figure. 3.**
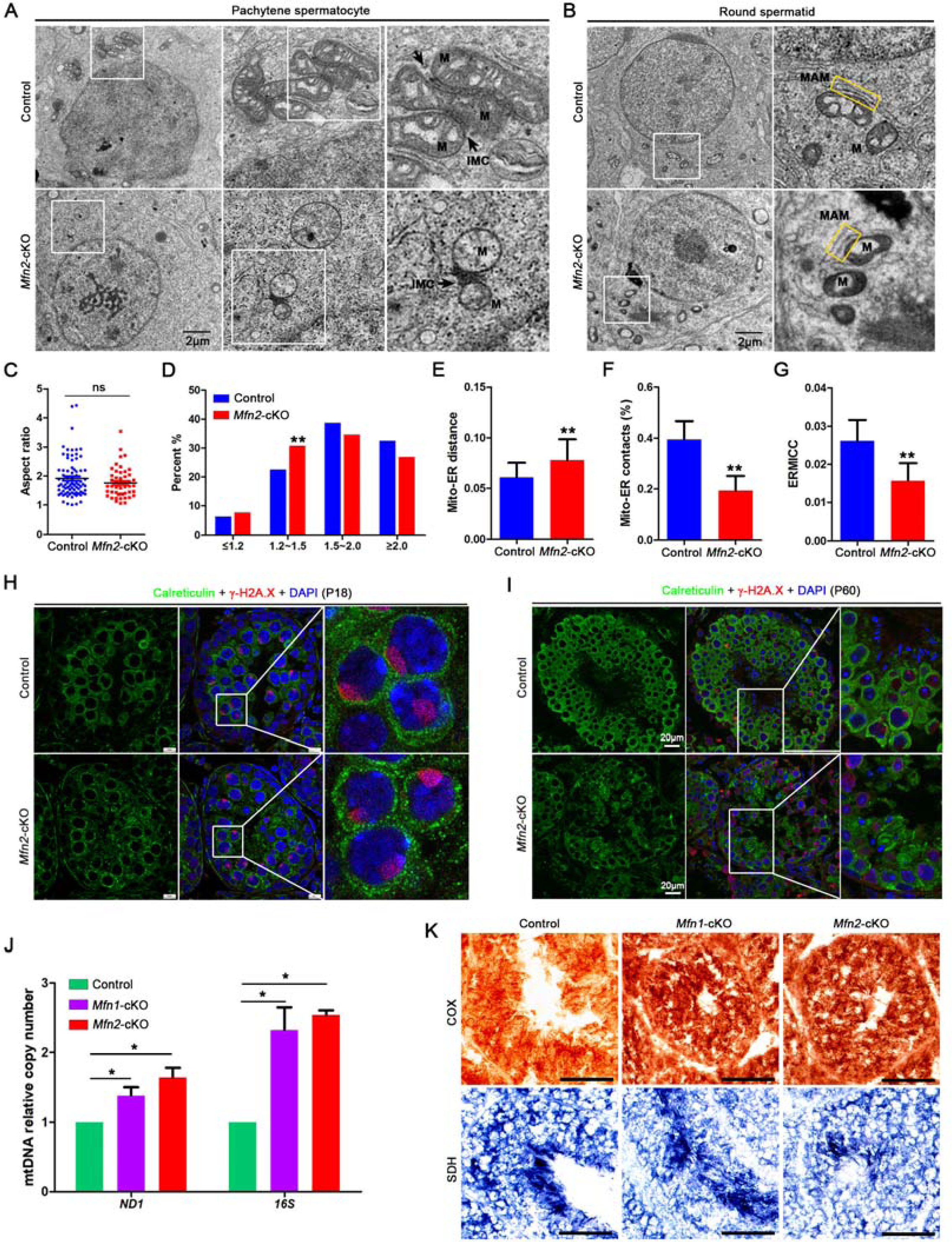
Loss of MFN2 in postnatal germ cells causes mitochondria, ER and MAM defects in testes. (**A**) TEM showing the fragmented and swelling mitochondria (M) in pachytene spermatocytes from *Mfn2*-cKO mouse testes at P60. Right panels represent the zoomed-in mitochondria from the square areas in the left panels. Arrows indicate inter-mitochondrial cement (IMC). (**B**) TEM showing the condensed swelling mitochondria without obvious cristae, ER fragmentation, and enlarged mitochondria-ER distance in round spermatid from *Mfn2*-cKO mouse testes at P60. Right panels represent the zoomed-in from the square areas in the left panels. Yellow rectangles indicate MAM structures. M, Mitochondria; ER, Endoplasmic reticulum; MAM, Mitochondria associated ER membranes. (**C**) Scatter plot showing the aspect ratio (AR) in round spermatids from control and *Mfn2*-cKO mouse testes at P60. AR represents the ratio of the major axis to the minor axis of mitochondria. (**D**) The distribution of AR in (C) showing an increased percentage of short mitochondria in *Mfn2*-cKO round spermatids. \*\**P*<0.01. (**E**) Histograms showing the distance between mitochondria and ER in round spermatids from control and *Mfn2*-cKO mouse testes at P60. \*\**P*<0.01. (**F**) Histograms showing the decreased contact area between mitochondria and ER in *Mfn2*-cKO round spermatids. \*\**P*<0.01. (**G**) Histograms showing the ERMICC values in round spermatids from control and *Mfn2*-cKO mouse testes at P60. ERMICC = mitochondrial-ER interface length / (mitochondrial perimeter x mitochondrial-ER distance). \*\**P* < 0.01. (**H**) Co-immunostaining of Calreticulin (an ER and MAM marker) and γ-H2A.X in testis sections from control and *Mfn2*-cKO mice at P18. Nuclei were stained with DAPI. Scale bar=10µm. (**I**) Co-immunostaining of Calreticulin and γ-H2A.X in testis sections from control and *Mfn2*-cKO mice at P60. Nuclei were stained with DAPI. Scale bar=20µm. (**J**) qPCR analyses showing the mtDNA copy numbers in control, *Mfn1*-cKO, and *Mfn2*-cKO mouse testes at P60. ND1 and 16S are the two genes encoded by mtDNA. \**P* < 0.05. (**K**) Respiratory enzymes COX (encoded by mitochondrial DNA) and SDH (encoded by nuclear DNA) staining of testis sections from adult control, *Mfn1*-cKO, and *Mfn2*-cKO mice. Scale bar=50µm.

Given that MFN2 supports structural and functional communication between mitochondria and ER, we next investigated whether the deletion of *Mfn2* in postnatal male germ cells disrupts MAM and ER homeostasis. In control testes, the majority of the mitochondria displayed close interactions with ER. However, the distance between mitochondria and ER increased by about 16%, and the percentage of mitochondria-ER contacts was significantly reduced by nearly 50% in *Mfn2*-cKO testes compared to that of controls (Figure.3E-F). Additionally, the ER-mitochondria contact coefficient (ERMICC) was also reduced by more than 30% in *Mfn2*-cKO testes compared with that of controls (Figure.3G). Notably, the ER displays tube-like cisternae structures in control spermatids, whereas it exhibits fragmentation in *Mfn2*-cKO spermatids (Figure.3B in yellow frame), implying that the ER structure disrupted upon MFN2 deletion in male germ cells. To further confirm these observations, calreticulin (a MAM and ER marker protein) immunostaining was employed to examine the MAM structure in P18 and P60 testes. Calreticulin displayed a diffused granular pattern in the cytoplasm of *Mfn2*-cKO spermatocytes at P18 instead of continuous perinuclear tubular localization exhibited in the controls (Figure.3H). Moreover, in P60 testes, the calreticulin signals in *Mfn2*-cKO germ cells appeared to be reduced and displayed diffused granular distribution compared to that of controls (Figure.3I). Of note, unlike the observation in *Mfn2*-cKO spermatocytes, calreticulin does not exhibit any diffused signal pattern in *Mfn1*-cKO spermatocytes (Supplementary Figure.S5B). Together, these data suggest MFN2 but not MFN1 may have a function in the regulation of the formation of MAM and ER structures.

Since dynamic mitochondrial fusion and fission are known processes of regulating mtDNA stability and energy production (Westermann, 2010; Palmer et al., 2011), we determined mtDNA copy number in *Mfn1*-cKO and/or *Mfn2*-cKO adult testes by RT-qPCR. Unexpectedly, we found that the mtDNA copy number appeared to increase in both *Mfn1*-cKO and *Mfn2*-cKO adult testes (Figure.3J), which is inconsistent with previous reports showing that conditional a single knockout of either *Mfn1* or *Mfn2* in skeletal muscle did not alter mtDNA copy number (Chen et al., 2010). The discrepancy and the heterogeneity of mitofusins imply a possible additional mechanism of MFN1/2 in regulating mtDNA stability between somatic and germ cells. Given that the physiological and ultrastructural evidence for mitochondrial dysfunction in *Mfn1*-cKO and/or *Mfn2*-cKO germ cells, we carried out the histochemical staining of cytochrome c oxidase (COX, complex IV, brown stain, encoded by mitochondrial DNA) and succinate dehydrogenase (SDH, complex II, blue stain, encoded by nuclear DNA) activity in adult testes to directly assess whether the function of respiratory complexes was affected. In line with the increased mtDNA copy number, COX activity increased in both *Mfn1*-cKO and *Mfn2*-cKO testis sections, whereas the SDH activity appeared unaltered (Figure.3K). These results show that the deletion of *Mfn1/2* in male germ cells leads to increased mitochondrial fission and dysregulated mitochondrial metabolism. Taken together, these data reveal that both MFN1 and MFN2 are essential for the maintenance of mitochondrial morphology in male germ cells, but MFN2, not MFN1 plays a critical role in maintaining MAM and ER structure integrity during male germ cell development.

### MFN2 interacts with Nuage-associated proteins and affects their expressions and piRNA production in testes

Since MFN2 displays a granular cytoplasmic localization in the cytoplasm of male germ cells (Figure.1), we assumed MFN2 maybe involve in mitochondria-associated germinal structure formation. To test this hypothesis, we selected four mitochondrial-associated germinal granule proteins (also terms of Nuage-associated proteins), including MIWI, DDX4, TDRKH, and GASZ, to perform the reciprocal immunoprecipitation (Co-IP) using MFN2 specific antibody in testis and cell line. We found that, in MFN2 antibody immunoprecipitants, MIWI, DDX4, GASZ, and TDRKH strongly detected in WT testes (Figure.4A). Likewise, in MIWI, DDX4, TDRKH, and GASZ antibody immunoprecipitants, MFN2 was detected in testes as well (Figure.4B-E). These results suggest that MFN2 interacts with MIWI, DDX4, TDRKH, and GASZ in testes. We also examined their interaction *in vitro* in the cell line. When the full length of MFN2 was ectopically co-expressed with MIWI, TDRKH, and DDX4 in HEK293T cells, MFN2 was pulled down by MIWI, TDRKH, and DDX4, indicating MFN2’s ability to interact with MIWI, TDRKH and DDX4 proteins (Figure.4F-G). Importantly, our previous studies have shown that all of these interacting proteins are within the MAM structures of mouse and human testes (Wang et al., 2018), suggesting MFN2 may function with the Nuage-associated proteins in maintaining MAM structure integrity.

**Figure. 4.**
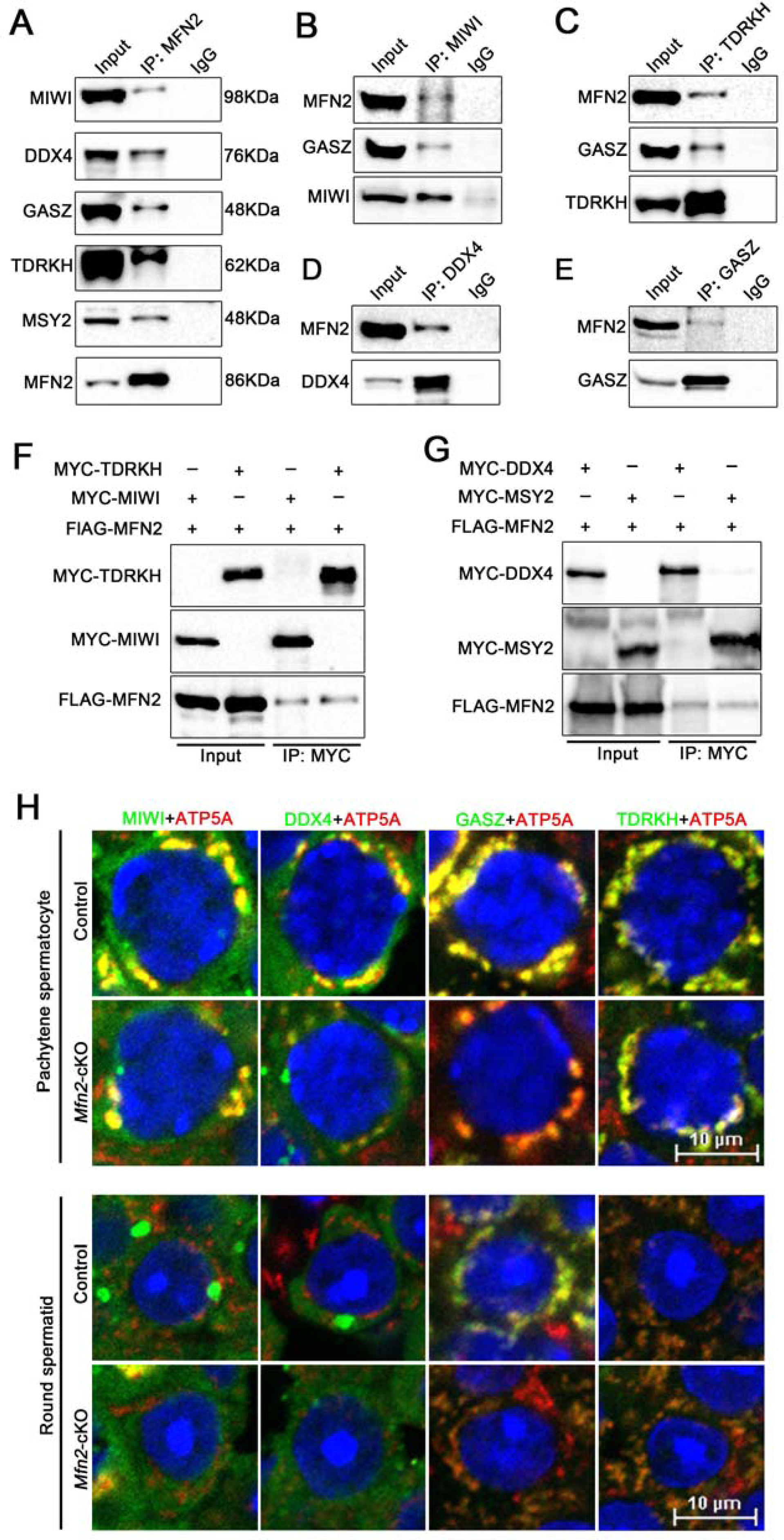
MFN2 interacts with nuage-associated proteins and regulates their expressions in spermatogenic cells. (**A**) Co-immunoprecipitation of MFN2 followed by western blot detection of MIWI, DDX4, GASZ, TDRKH, and MSY2 in adult mouse testes. (**B-E**) Reciprocal immunoprecipitation showing that MFN2 detected in the immunoprecipitants from MIWI (b), TDRKH (c), DDX4 (d), and GASZ (e) in the adult mouse testes, respectively. (**F**) MFN2 interacts with MIWI and TDRKH *in vitro*. HEK293T cells were transfected with indicated plasmids. After 48h of transfection, immunoprecipitation was performed using anti-MYC antibodies and detected by anti-MYC and anti-FLAG antibodies, respectively. (**G**) MFN2 interacts with DDX4 and MSY2 *in vitro*. HEK293T cells were transfected with indicated plasmids. Immunoprecipitation was performed using anti-MYC antibody and detected by anti-MYC and anti-FLAG antibodies, respectively. (**H**) Co-immunostaining of ATP5A with MIWI, DDX4, GASZ, and TDRKH in pachytene spermatocyte and round spermatid from control and *Mfn2*-cKO mice at P60, respectively. Scale bar=10 µm.

We next examined whether the expression levels of MIWI, DDX4, GASZ, and TDRKH interacting proteins were affected upon deletion of MFN2 in testes. Co-immunostaining of *Mfn2*-cKO and control testes revealed ectopic expression of MIWI, DDX4, and GASZ in *Mfn2*-cKO testes in IMC and CB structures while TDRKH displayed no apparent changes (Figure.4H and Supplementary Figure. S5C-E). In *Mfn2*-cKO testes, both MIWI and DDX4 showed weaker staining in the IMC of pachytene spermatocytes and barely detected in the CB of round spermatids (Figure.4H and Supplementary Figure.S5C-D). Of note, unlike the perinuclear granular mitochondrial localization showed in the control spermatocytes, DDX4 displayed a diffuse cytoplasmic expression pattern in *Mfn2*-cKO spermatocytes (Supplementary Figure.S5D). GASZ also showed a decreased signal in both pachytene spermatocytes and round spermatids within *Mfn2*-cKO testicular sections (Figure.4H and Supplementary Figure.S5E). Since MFN1 was reported to interact with GASZ and is essential for spermatogenesis(Zhang et al., 2016), we also compared the expression and localization of DDX4, GASZ, and TDRKH in *Mfn1*-cKO adult testes. However, unlike *Mfn2*-cKO testes, we found no obvious changes in expression or localization in *Mfn1*-cKO testes (Supplementary Figure.S5F-H).

Given the Nuage-associated proteins are documented in regulating the piRNA biogenesis pathway, which prompts us to determine whether MFN2 participates in the piRNA pathway. Due to the morphology and the cell populations are comparable at P25 between WT and *Mfn2*-cKO testes, we chose the P25 testes to examine the abundance and size of piRNA distributions through small RNA sequencing. The results showed about a 50% decrease of the whole piRNA populations after normalized with miRNA counts (Supplementary Figure.S6A and Table S1), suggesting that MFN2 might take part in piRNA biogenesis through its interaction with the Nuage proteins. We next tried to ask whether the retrotransposons were affected in *Mfn2*-cKO testes because the Nuage-associated proteins have reported repressing retrotransposons for maintaining the genome integrity of male germline. By RT-qPCR assays, we analyzed the mRNA expression levels of retrotransposons, including LINE1 and IAP, and found that there are no significant changes between WT and *Mfn2*-cKO testes (Supplementary Figure.S6B). Consistent with the mRNA level of LINE1, the LINE1 ORF1 protein expression showed no activation both in P25 (data not shown) and P60 *Mfn2*-cKO testes (Supplementary Figure.S6C). Taken together, these results indicate that MFN2 plays a crucial role in the formation and/or maintenance of the network of RNA processing proteins and piRNA pathways in the Nuage during spermatogenesis.

### RNA-seq reveals differential expression genes involved in mRNA processing in spermatogenic cells

To understand the underlying molecular mechanism of MFN2 in regulating spermatogenic cell development, we performed whole-transcriptome sequencing to assess the differential expression genes in purified pachytene spermatocytes (PS) and round spermatids (RS) from WT and *Mfn2*-cKO adult testes. After RNA deep sequencing, we performed pairwise differential gene expression comparisons to define the genes whose expression changed by at least 2-fold (*P* ≤ 0.05) when WT and *Mfn2*-cKO groups were compared.

Hierarchical clustering of differentially expressed genes (DEG) showed that, in pachytene spermatocytes, a total of 4046 genes up-regulated and 5324 genes were down-regulated in *Mfn2*-cKO mice compared to that of WT mice. In round spermatids, a total of 3756 genes up-regulated and 3186 genes were down-regulated in *Mfn2*-cKO mice compared to that of WT mice (Figure.5A-B, Supplementary Fig.7SA-B and Table S2-3). Principal component analysis (PCA) separated the data into four sub-groups clustering each separately as WT control and *Mfn2*-cKO pachytene spermatocytes and round spermatids, showing the gross changes of the transcriptome between WT and *Mfn2*-cKO transcriptome (Supplementary Figure.S7C).

**Figure. 5.**
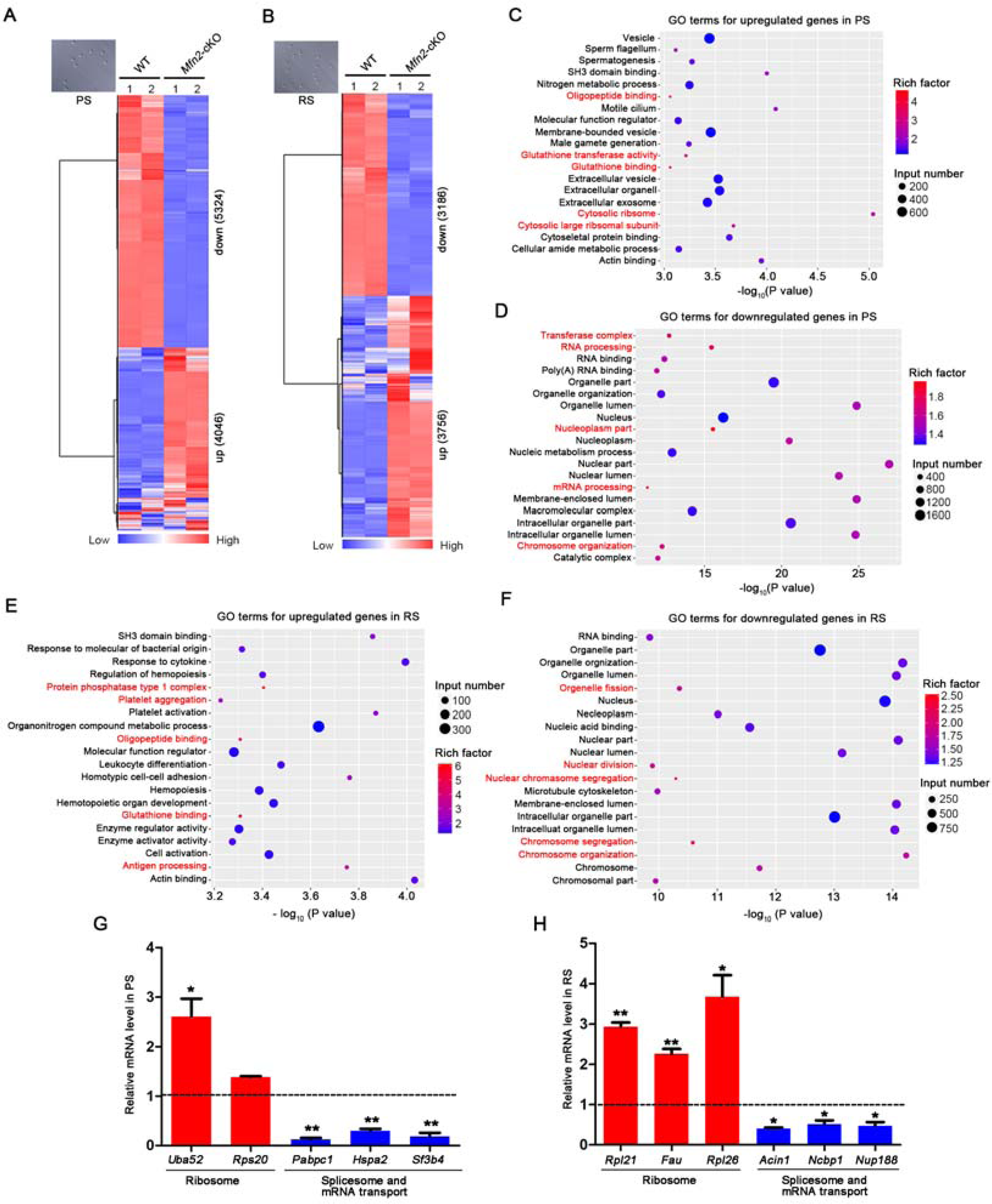
Global gene expression altered in *Mfn2*-cKO spermatogenic cells. (**A-B**) Heatmap is showing the differential expression genes in purified pachytene spermatocytes (A) and round spermatids (B) from adult WT and *Mfn2*-cKO testes, respectively. The upper left images are showing the purity and morphology of isolated pachytene spermatocytes (PS) and round spermatids (RS), respectively. Significantly regulated genes have a *P*-value of <0.05 and a fold change of > 2.0. Two biological replicates indicated in the heat-map. (**C-D**) Gene ontology (GO) term analyses showing the top 20 GO terms for the up-regulated (C) and down-regulated (D) genes in *Mfn2*-cKO pachytene spermatocytes. Rich factor= (number of differentially expressed genes in GO term)/(total number of genes in GO term). A larger rich factor indicates higher enrichment. Top 5 GO terms by rich factor highlighted in red. (**E-F**) Gene ontology (GO) term analyses showing the top 20 GO terms for the up-regulated (E) and down-regulated (F) genes in *Mfn2*-cKO round spermatids. Top 5 GO terms by rich factor highlighted in red. (**G-H**) Validation of the up- and down-regulated genes selected from RNA-seq data in purified pachytene spermatocytes (G) and round spermatids (H) by RT-qPCR analyses.

Gene Ontology (GO) was then applied to assess what molecular functions and biological processes altered upon a lack of *Mfn2* in testes. In pachytene spermatocytes, the most up-regulated genes in *Mfn2*-cKO mice mainly involved regulation of biological quality and localization (Supplementary Figure.S7D). The down-regulated genes were primarily associated with the metabolic process (Supplementary Figure.S7E). Similarly, in round spermatids, the up-regulated genes in *Mfn2*-cKO mice were mostly distributed in vesicle fraction and associated with those that regulate phosphorus metabolic processes, and the down-regulated genes were distributed in cell and intracellular part and enriched in the cellular and metabolic processes (Supplementary Figure.S7F-G). Importantly, when the top 20 GO terms analyzed by organized *P*-value and rich factor, we found that the most up-regulated genes in *Mfn2*-cKO pachytene spermatocytes were mainly involved in glutathione binding, oligopeptide binding, glutathione transferase activity, cytosolic large ribosomal subunit and cytosolic ribosome, which run the top 5 enrichment sorted by rich factor. Besides, sperm flagellum, spermatogenesis and motile cilium were included in the top 20 GO terms (Figure.5C). While for the down-regulated genes, mRNA processing, nucleoplasm part, RNA processing, transferase complex and chromosome organization harbored the higher rich factor (Figure.5D). By contrast, in *Mfn2*-cKO round spermatids, the top 5 enrichment sorted by rich factor for the up-regulated genes were involved in protein phosphatase type 1 complex, glutathione binding, oligopeptide binding, antigen processing and platelet aggregation (Figure.5E), and for down-regulated genes with higher rich factor were involved in nuclear chromosome segregation, chromosome segregation, nuclear division, organelle fission and chromosome organization (Figure.5F).

KEGG pathway analysis revealed that, in pachytene spermatocytes, the top 5 pathways by *P-*value associated with up-regulated genes belonged to glycolysis/glucogenesis, ribosome, HIF-1 signaling, biogenesis of amino acids and carbon metabolism, and the top 5 pathways involved in down-regulated genes were related to spliceosome, RNA transport, Huntington’s disease, ribosome biogenesis in eukaryotes and cell cycle (Supplementary Figure.S7H-I). In round spermatids, the KEGG pathways associated with up- and down-regulated genes were similar to that of in pachytene spermatocytes, as emphasized for the top 5 pathways (Supplementary Figure.S7J-K).

To validate the RNA-seq data, we randomly selected several genes associated with ribosome, splicesome and mRNA transport pathways for RT-qPCR analysis. In line with the RNA-seq data, genes involved with ribosome transport pathways up-regulated, whereas genes associated with splicesome and mRNA transport pathways were down-regulated in both pachytene spermatocytes and round spermatids (Figure.5G-H). Taken together, these RNA-Seq analyses support the view of an association of MFN2 with cytosolic ribosomes, the spliceosome and mRNA processing pathways essential for male germ cell development act in an integral manner.

### MFN2 ablation results in misregulated alternative splicing events

Given that our RNA-seq data suggest MFN2 associates with mRNA processing, we next asked whether alternative splicing (AS) events were affected in spermatogenesis. To test this possibility, we performed alternative transcript splicing analyses for the RNA-seq data on several of regulated splicing events in both *Mfn2*-cKO pachytene spermatocytes and round spermatids. Indeed, a total of 1604 misregulated splicing events were identified in 1001 genes in *Mfn2*-cKO pachytene spermatocytes, and 986 misregulated splicing events in 684 genes in *Mfn2*-cKO round spermatids were found (Figure.6A-B and Supplementary Table S4-5). Interestingly, the up- and down-regulated events account for ∼50% of the total changed events separately in both pachytene spermatocytes and round spermatids (Figure.6A-B).

**Figure. 6.**
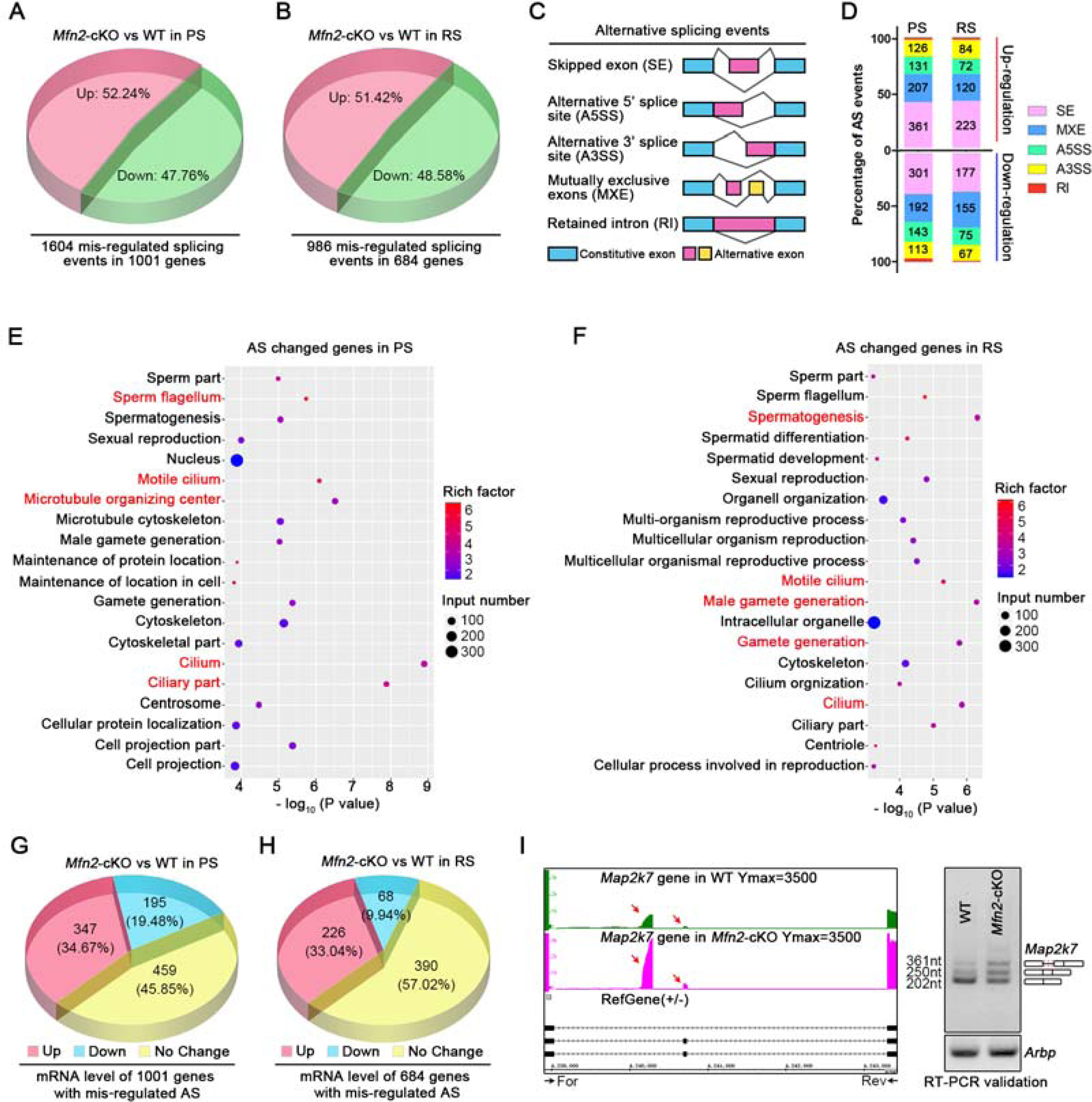
Ablation of MFN2 in postnatal germ cells leads to misregulated alternative splicing (AS) in spermatogenic cells. (**A-B**) Pie charts showing the percentages of misregulated splicing events identified from RNA-seq data in pachytene spermatocytes (A) and round spermatids (B) from WT and *Mfn2*-cKO mice (fold change> 2; *P* < 0.05). (**C**) Different types of alternative splicing are shown. In these graphics, boxes represent exons and lines represent introns. Exon regions included in the message by alternative splicing are colored with pink and yellow, while constitutive exons are shown in blue. (**D**) Stacked bar plot showing the ratio of AS status including up-(upper panel) and down-regulated (lower panel) events in pachytene spermatocytes (PS) and round spermatids (RS). Numbers on the bars are the amount of corresponding misregulated splicing events. (**E-F**) Gene ontology (GO) term analyses showing the top 20 GO terms for the genes with altered splicing events in pachytene spermatocytes (E) and round spermatids (F). Top 5 GO terms by rich factor selection highlighted in red. (**G-H**) Pie charts showing the percentages of mRNA expression level alterations in 1001 genes with misregulated alternative splicing in pachytene spermatocytes (G), and 684 genes with misregulated alternative splicing in round spermatids (H). (**I**) Screenshot from the IGB (Integrated Genome Browser) software showing high inclusion levels for an intron in the *Map2k7* gene in *Mfn2*-cKO pachytene spermatocytes (the left panel), and RT-PCR validation of the misregulated alternative splicing in WT and *Mfn2*-cKO pachytene spermatocytes (right panel). *Arbp* served as an internal control.

We then further analyzed the distribution of five typical AS types for those misregulated splicing events in pachytene spermatocytes and round spermatids, including skipped exon (SE), alternative 5’ splice site (A5SS), alternative 3’ splice site (A3SS), mutually exclusive exons (MXE) and retained intron (RI). The percentage of the five AS events analyses showed a similar distribution ratio with the SE highest and RI lowest in both pachytene spermatocytes and round spermatids (Figure.6C-D). Strikingly, GO analyses showed that the genes with abnormal splicing events enriched in spermatogenesis and male gamete generation in both pachytene spermatocytes and round spermatids (Figure.6E-F). Importantly, in pachytene spermatocytes, the top 5 enriched GO terms were cilium, ciliary part, microtubule organizing center, motile cilium and sperm flagellum (Figure.6E).

In round spermatids, the top 5 enriched GO terms were spermatogenesis, male gamete generation, cilium, gamete generation and motile cilium (Figure.6F). Thereinto, among these genes with abnormal alternative splicing, ∼34.67% of genes displayed up-regulated, and 45.85% genes showed down-regulated transcriptional mRNA levels in pachytene spermatocytes. Additionally, ∼33.04% genes with up-regulated and 57.02% genes with down-regulated transcriptional levels observed in round spermatids (Figure.6G-H). To further verify the modulation of MFN2 in alternative splicing, we selected a mis-spliced gene, *Map2k7*, with intron retention in both *Mfn2*-cKO pachytene spermatocytes and round spermatids based on the RNA-seq analysis as a validated gene by semi-quantitative PCR with specific primers. *Map2k7* encodes a protein kinase highly expressed in testis and acts as an essential component of the MAPK signal transduction pathway to regulate mitochondrial death signaling pathway and apoptosis(Yao et al., 1997; Tournier et al., 2001). We successfully verified the aberrant splicing pattern of *Map2k7* gene in *Mfn2*-cKO testes (Figure.6I). Taken together, these data indicate that both the transcriptional levels and alternative splicing processes of genes involved in spermatogenesis were affected upon MFN2 ablation in germ cells.

### MFN2 associates with mRNA translational machinery to regulate mRNA fates in testes

Because some of the MFN2 interaction Nuage-associated proteins, like MIWI and DDX4, are associated with RNA processing and inactive translational mRNAs in spermatogenesis(Grivna et al., 2006; Nagamori et al., 2011), we wanted to determine whether MFN2 plays a role in translation. To test this relationship, we first subjected adult testicular extracts to sucrose density gradient fractionation (Figure.7A). MFN2 is co-sedimented with both monosome (80S) and polysome fractions (Figure.7A). Similar to RPS6, the addition of EDTA to dissociate the large and small ribosomal subunits shifted MFN2 to the ribonucleoprotein (RNP) fractions of the gradient. Combined with the RNA-Seq data, these results suggest that MFN2 could associate with cytoplasmic monosomes and polysomes. Thus, we subsequently evaluated whether MFN2 regulates global mRNA translation in spermatogenic cells through ribosome profiling assays in control and *Mfn2*-cKO testes. Interestingly, *Mfn2*-cKO testes displayed a similar polysome proportion as the control, yet an increased ratio of polysomes to monomers (80S)(Wrobel et al., 2015) (Figure.7B-C). This result indicates the activation of global mRNA translation upon loss of MFN2 in testes.

**Figure. 7.**
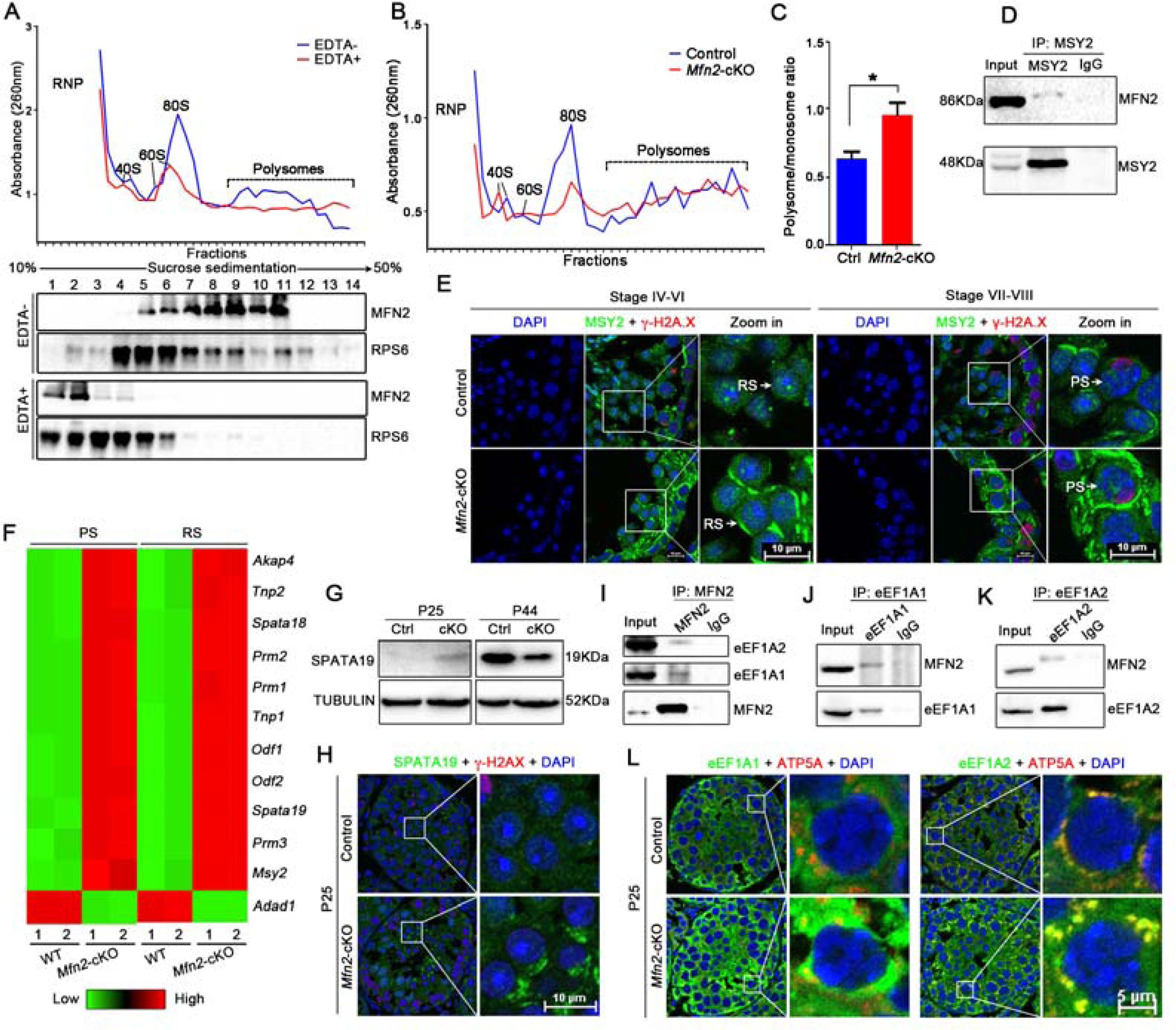
MFN2 associates with mRNA translation processes and regulates gamete-specific mRNA fates. (**A**) Untreated (EDTA-) and EDTA-treated (EDTA+) post-nuclear testicular extract from adult WT mice were fractionated on linear 10–50% sucrose density gradients and analyzed by UV spectrometry. The distribution of MFN2 protein is detected by Western blotting. RPS6 protein serves as a positive control. (**B**) Cytoplasmic polysome profiling of control (blue line) and *Mfn2*-cKO (red line) testes at P35. (**C**) Quantifications of the ratio of polysomes to monosomes (80S) in control (Ctrl) and *Mfn2*-cKO testes are shown. (**D**) Immunoprecipitation showing MFN2 was detected in the immunoprecipitants from MSY2 in the adult mouse testes. (**E**) Co-immunostainings of MSY2 and γ-H2A.X in stage IV-VI and VII-VIII seminiferous tubules from adult control and *Mfn2*-cKO testes. Nuclei were stained with DAPI. RS, round spermatids; PS, pachytene spermatocytes. Scale bar=10µm. (**F**) Heat-map is showing the differential expression levels of 12 MSY2-bound male gamete-specific mRNAs in both pachytene spermatocytes (PS) and round spermatids (RS) from RNA-Seq data in WT and *Mfn2*-cKO mice. (**G**) Western blots analysis of the protein levels of *Spata19* in control and *Mfn2*-cKO testes at P25 and P44, respectively. (**H**) Co-immunostaining of SPATA19 and γ-H2A.X in P25 testes from control and *Mfn2*-cKO mice. Nuclei were stained with DAPI. Scale bar=10µm. (**I**) Reciprocal immunoprecipitation showing eEF1A2 and eEF1A1 were detected in the immunoprecipitants from MFN2 in the adult mouse testes. (**J-K**) Reciprocal immunoprecipitation showing MFN2 was detected in the immunoprecipitants from eEF1A1 (J) and eEF1A2 (K) in the adult mouse testes, respectively. (**L**) Co-immunostainings of eEF1A1 and ATP5A (left panel), eEF1A2 and ATP5A (right panel) in P25 testes from control and *Mfn2*-cKO mice. Nuclei were stained with DAPI. Scale bar=10µm.

Since MSY2, as a germ cell-specific DNA/RNA-binding protein, exclusively enriched in both the meiotic and post-meiotic germ cells (Gu et al., 1998), and reported to participate in both transcription and mRNA translation (Yang et al., 2005b). Considering the above, we next asked whether MFN2 complexes with MSY2 to regulate mRNA translation in testes. Co-IP experiments revealed that MFN2 could interact with MSY2 both *in vivo* and *in vitro* (Figure.4A and G; Figure.7D). Because our unbiased RNA-seq data showed an increased *Msy2* mRNA level in both *Mfn2*-cKO pachytene spermatocytes and round spermatids, we then wanted to examine the protein level of MFN2 in adult control and *Mfn2*-cKO testes by immunofluorescence. In control testes, MSY2 was broadly expressed in nearly all types of spermatogenic cells and showed a disperse distribution in the cytoplasm and nucleus in pachytene spermatocytes and round spermatids. However, in *Mfn2*-cKO testes, the cytoplasmic signals of MSY2 increased in pachytene spermatocytes and round spermatids, and the MSY2 foci signals within the round spermatid nucleus had almost disappeared (Figure.7E). Considering that MSY2 marks specific mRNAs in the nucleus for cytoplasmic storage and links transcription and mRNA storage/translational delay in meiotic and post-meiotic male germ cells (Yang et al., 2005a), we analyzed the transcriptional levels of MSY2-bound mRNAs in our RNA-seq data. Intriguingly, most of the MSY2-bound gamete-specific mRNAs appeared to upregulate in *Mfn2*-cKO pachytene spermatocytes and round spermatids, including *Msy2* itself (Figure.7F and Supplementary Table S6). To further explore the role of MFN2 in translation, we examined if any proteins mistranslated from the MSY2-bound gamete-specific mRNAs. Since SPATA19 (also known as SPERGEN1) is a mitochondrial protein previously reported to express in haploid spermatids at P28 testes(Doiguchi et al., 2002; Matsuoka et al., 2004) and germ cell-specific *Spata19* knockout mice displayed male sterility(Mi et al., 2015), we thus chose SPATA19 to examine the protein level of *Spata19* in P25 testes from control and *Mfn2*-cKO mice. Notably, we found that the protein of SPATA19 can be detected in P25 *Mfn2*-cKO round spermatids by Western blot or immunofluorescence, whereas undetectable in control round spermatids (Figure.7G-H). These data show that *Spata19* was early activated and translated when a loss of MFN2 in male germ cells, suggesting MFN2 could regulate mRNA translation.

To determine how MFN2 regulates mRNA translation, we examined the interplay of MFN2 with the eukaryotic elongation factors (eEF1A1 and eEF1A2) in testes, which are critical regulators during translational processes (Negrutskii and El’skaya, 1998; Sasikumar et al., 2012). As expected and revealed by reciprocal Co-IP, both eEF1A1 and eEF1A2 could interact with MFN2 (Figure.7I-K). Moreover, immunofluorescence showed that the signals of eEF1A1 and eEF1A2 increased in *Mfn2*-cKO pachytene spermatocytes. Of note, we found that both eEF1A1 and eEF1A2 appeared to express in the cytoplasm of pachytene spermatocytes, and eEF1A2 co-localized with ATP5A (a mitochondria marker) but not eEF1A1 (Figure.7L). These data suggest that the early activated mRNA translation in Mfn2-cKO testes, especially for some gamete-specific mRNAs, is likely due to the disassociation of the interplay of MFN2 with elongation factor eEF1A1 and eEF1A2. Taken together, these results indicate that *Mfn2* is associated with mRNA translational machinery by interaction with several Nuage-associated proteins (like MIWI) and control the fates of gamete-specific genes during spermatogenesis.

## Discussion

Mitofusins, MFN1 and MFN2, are the first known mediators of mitochondrial fusion proteins in mammals (Hales and Fuller, 1997). Their functions in neuron and heart have well explored(Dietrich et al., 2013; Schneeberger et al., 2013; Hall et al., 2016; Jiang et al., 2018), yet their roles in spermatogenesis are poorly understood. In this study, we showed that the germ cell-specific deletion of *Mfn2* disrupts the mitochondria, ER and MAM structure, gamete-specific mRNA translation processes, resulting in male sterility. Importantly, the phenotype of *Mfn2*-cKO mice is age-dependent, with germ cells beginning their decline after P35. With increasing age, vacuolation of the seminiferous tubule becomes severe. In comparison, the *Mfn1*-cKO shows vacuolization as early as P28 in this study. Interestingly, *Vasa*-Cre mediated *Mfn1* knock out mice (*Vasa*-Cre; *Mfn1*^flox/flox^) were also infertile and displayed very severe phenotypes showing seminiferous tubule vacuolization as early as P14(Zhang et al., 2016). The discrepancy of phenotypes between our *Mfn1*-cKO mice (*Stra8*-Cre; *Mfn1*^flox/flox^) and the reported mice (*Vasa*-Cre; *Mfn1*^flox/flox^) possibly reflects the differences in Cre recombinase expression since *Vasa*-Cre appears much earlier than *Stra8*-Cre in the male germ. Remarkably, the *Mfn1*/*2* double knockout testes showed a more severe phenotype of near loss of pachytene spermatocytes and round spermatids, and the accumulation of mitochondria to one side of germ cell cytoplasm.

Despite some of the MFN2 interaction proteins showed abnormal expression in *Mfn2*-cKO testes, including GASZ, MIWI, and DDX4, they didn’t show an apparent change in *Mfn1*-cKO testes. This data is in agreement with the previous report in which *Vasa*-Cre mediated *Mfn1* conditional knockout mice showed no apparent alteration in the localization of GASZ and DDX4 (Zhang et al., 2016). Therefore, previous *Mfn1* or *Mfn2* genetic mutation functional studies in Purkinje cells and oocytes require much careful consideration (Chen et al., 2007; Hou et al., 2019), as their interaction and role in different mammalian cell development may differ. In this study, we found that both MFN2 and MFN1 are required for spermatogenesis and male fertility, but they may exhibit a different mechanism to control spermatogenesis during male germ cell development. Indeed, our study revealed that MFN2 might work with MFN1 coordinately in spermatogenesis by taking part in different molecular pathways.

Despite a myriad of published literature focused on the function of MAM structure in neuronal systems and metabolism with several human diseases(Paillusson et al., 2016), the role of MAM structure in spermatogenesis has not reported yet. Our previous work observed that the abundance of MAM in spermatogenic cells, and many proteins identified in mass spectroscopy data were crucial regulators of spermatogenesis (Wang et al., 2018). Unexpectedly, several MFN2 interaction proteins identified in this work, including MIWI, DDX4, GASZ, and TDRKH, were present in the MAM fraction from both mouse and human testes. The data presented in the current study showed that in *Mfn2*-cKO, pachytene spermatocytes and round spermatids, MIWI, DDX4, and GASZ expression decreased. As part of the testicular MAM structure, it is likely that MFN2 and its interacting proteins serve as MAM proteins and form a protein complex or a scaffold to regulate spermatogenesis. This complex is a dynamic partner regulated by the mitochondria fusion and fission process. Once MFN2 deleted, the morphology of mitochondria, ER and MAM structures of germ cells disrupted. Therefore, the current study, for the first time, provides a possible new role of MFN2 as the MAM component in spermatogenesis and spermatogenic cell development. Specifically, the proteins located in MAMs are possibly responsible for regulating the piRNA biogenesis and mRNA translational pathways as they are a feature of Nuage-associated proteins in germ cells. The fact that we observed the aberrant piRNA production in *Mfn2*-cKO testes, suggesting MFN2 might have a function in piRNA biogenesis. However, we could not exclude the possibility in which caused by mislocalized Nuage-associated proteins, such as MIWI, DDX4, GASZ, and TDRKH, et al., in *Mfn2*-cKO mice. It is worth noting that although the thickness of the IMC structure appeared to increase in *Mfn2*-cKO pachytene spermatocytes, this likely reflects an increased distance between mitochondria. Therefore, it indicated that the formation of IMC might is independent of MFN2 and possibly caused by misregulated Nuage-associated proteins in *Mfn2*-cKO pachytene spermatocytes as well.

An exciting finding in our study is that MFN2 enriched in monosome and polysome fractions in spermatogenic cells and cooperates with MSY2 to mediate MSY2-bound gamete-specific mRNA translation. MSY2 showed increased cytosolic expression in both pachytene spermatocytes and round spermatids upon MFN2 depletion. This discovery raises the possibility that MFN2 partners with MSY2-bound mRNA for its function because MSY2 was enriched in RNP fraction and regulates mRNA stability, storage and translation delay in spermatogenesis (Yang et al., 2005a). Additionally, we identified, in this study, several MFN2 interaction partners, such as MIWI and DDX4, localized in the Nuage (IMC and CB) and associated with translational machinery. MIWI enriched in both RNP and polysome fractions and associates with both mRNAs and piRNAs in cytosolic ribonucleoprotein and polysomal fractions (Grivna et al., 2006), whereas DDX4 has reported stimulating translation initiation of specific mRNAs through interaction with the general translation initiation factor eIF5B(Johnstone and Lasko, 2004; Liu et al., 2009). Such partnerships might be essential for initiating spermiogenesis and gamete-specific mRNA translational delay because spermatogenesis in both *Miwi* and *Msy2* mutants are all arrested at the post-meiotic stage (Deng and Lin, 2002; Yang et al., 2005b).

In mammalian cells, most protein-coding genes are disrupted by intervening sequences (introns), which need to precisely and effectively removed through alternative splicing in different cell types or tissues (Baralle and Giudice, 2017). Interestingly, the mammalian testis is among the tissues with the highest transcriptome complexity, including alternative splicing regulation, especially in the spermatocyte and spermatids (Ramskold et al., 2009; Soumillon et al., 2013; Baralle and Giudice, 2017; Estill et al., 2019). In this study, we revealed over a thousand misregulated splicing events occur in *Mfn2*-cKO spermatogenic cells by RNA-seq analyses. Besides, most genes identified with misregulated splicing events in this study were critical for spermatogenesis and gamete generation. However, it is worthy to point out that alternative splicing usually occurs in the nucleus or adjacent nuclear envelope, but MFN2 mainly localized in the cytoplasm of germ cells. Therefore, the misregulation of alternative splicing observed in *Mfn2*-cKO germ cells were surmised to be secondary defects caused by *Mfn2* deletion. The possible explanation is that disruption of an RNA binding proteins/mRNA translation protein complex upon *Mfn2* mutated in testes, especially MSY2, which reported to bind mRNA directly and regulate their storage and translation, would reconcile these observations.

In conclusion, we report, in this study, a novel role of MFN2 in RNA processing through interactions with Nuage-associated proteins and mRNA translation proteins in male germ cells during spermatogenesis. We propose that MFN2 associates with several mRNA translation associated proteins such as MSY2, MIWI, and DDX4 at the cytoplasmic Nuage and/or MAM, and it likely serves as a scaffold to recruit mRNA, polysomes and other RNA binding proteins in both pachytene spermatocytes and round spermatids, which further work together with the elongation factor1 alpha subunits (like eEF1A1 and eEF1A2) to control gamete-specific transcripts delay during spermatogenesis (Figure.8A-B). These are beyond its known functions in regulating mitochondrial fusion processes. Furthermore, our research suggests that Mitofusins (both MFN1 and MFN2) are essential for mouse spermatogenesis and male fertility, and provide the first evidence for mitofusin proteins associated with RNA processing and translational machinery in regulating germ cell development during mouse spermatogenesis.

**Figure. 8.**
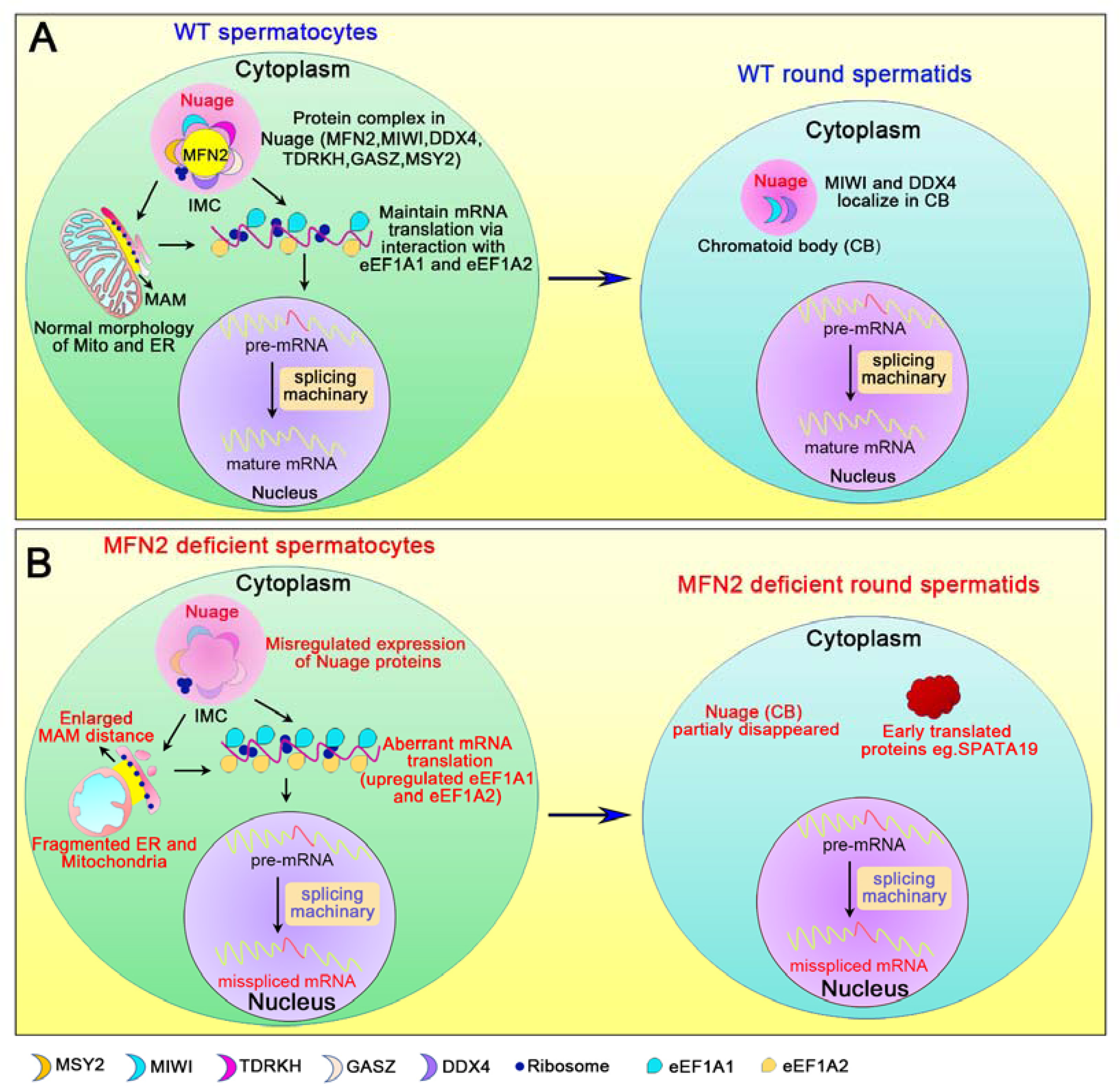
The working model illustrates the function of MFN2 in male germ cell development. (A) In WT spermatocytes, MFN2 cooperates with Nuage associated proteins in pachytene spermatocytes and round spermatids, including MIWI, DDX4, TDRKH, GASZ, and MSY2 to form interaction complexes for maintaining normal morphology of mitochondria, MAM and ER, and mRNA translation via interaction with the eukaryotic elongation factor eEF1A1 and eEF1A2. **(B)** In MFN2 deficient pachytene spermatocytes and round spermatids, Nuage-associated proteins are ectopically expressed, including a decrease of MIWI, DDX4, and GASZ expression, and an increase of MSY2 expression. Meanwhile, in the cytoplasm, the disassociated MFN2-Nuage protein complexes caused some proteins are misregulated in mRNA and/or protein level, especially gamete-specific genes, such as SPATA19 is early translated, which leads to disrupted male germ cell development and male fertility.

## Materials and methods

### Animals and ethics statement

All animal procedures were approved by the Institutional Animal Care and Use Committee (IACUC) of Tongji Medical College, Huazhong University of Science and Technology, and the mice housed in the specific pathogen-free facility of Huazhong University of Science and Technology. All experiments with mice were conducted ethically according to the Guide for the Care and Use of Laboratory Animal guidelines. Floxed *Mfn1* or *Mfn2* mice were created previously(Chen et al., 2007). The mice were harboring the floxed *Mfn1* or

*Mfn2* allele purchased from the Jackson Laboratory (Stock no. 026401 and 026525). *Mfn1*^flox/flox^ and/or *Mfn2*^flox/flox^ female mice were crossed with *Stra8*-Cre male mice (*Stra8*-Cre mice was purchased from the Jackson Laboratory, stock no. 008208) to obtain *Stra8*-Cre; *Mfn1*^+/flox^ and/or *Mfn2*^+/flox^ males, then the *Stra8*-Cre; *Mfn1*^+/flox^ and/or *Mfn2*^+/flox^ male mice were further bred with *Mfn1*^flox/flox^ and/or *Mfn2*^flox/flox^ to obtain *Stra8*-Cre; *Mfn1*^flox/del^ and/or *Mfn2*^flox/del^ (designated as *Mfn1*-cKO and/or *Mfn2*-cKO) males.

### Antibodies

The details of all commercial antibodies used in this study are presented in Supplementary Table S7.

### Histological analysis, immunostaining and imaging

Testes and epididymides from control and cKO mice were collected and fixed in Bouin’s solution (SIGMA, HT10132) at 4°C overnight and then washed with 75% alcohol 5 times for 30 minutes each time. Samples were then embedded in paraffin, and 5μm sections were cut and stained with periodic acid-Schiff (PAS) after being dewaxed and rehydrated. Slides were then mounted with Permount (Fisher Chemical^TM^SP15-100) and imaged with a ZEISS microscope. For immunofluorescence staining, testes were fixed in 4% PFA in PBS overnight at 4°C and embedded in the Tissue-Tek O.C.T. compound (Sakura Finetek, 4583) on dry ice. A 5μm thick section was cut and microwaved in 0.01M sodium citrate buffer (pH=6.0) for 5 min to retrieve antigen. After rinsing with PBS three times, the sections were blocked in blocking solution (containing 3% normal goat serum and 3% fetal bovine serum in 1% bovine serum albumin) for 1h at room temperature. Testis sections then incubated with primary antibodies (Supplementary Table S7) diluted in blocking solution overnight at 4°C. After washing with PBS, sections were incubated with Alexa Fluor 488 goat anti-rabbit IgG (1:500; A32731, Invitrogen) and/or Alexa Fluor 594 goat anti-mouse IgG (1:500, A11032, Invitrogen) for 1h at room temperature and then stained with DAPI for 5 min, washed in PBS and mounted using 80% glycerol. Confocal fluorescence microscopy was conducted using a confocal A1 laser microscope (Nikon, Japan).

### TUNEL analyses

Testes were fixed in Bouin’s solution, embedded in paraffin and sectioned (5μm). TUNEL staining performed using One Step TUNEL Apoptosis Assay Kit (Meilunbio Cat. MA0223) according to the manufacturer’s instructions. Images were obtained with a FluoView 1000 microscope (Olympus, Japan).

### Transmission electron microscopy (TEM)

TEM was performed as described previously with some modifications(Wang et al., 2018). In brief, testes from control and cKO were fixed in 0.1M cacodylate buffer (pH=7.4) containing 3% paraformaldehyde and 3% glutaraldehyde plus 0.2% picric acid for 2h at 4°C, then for 1h at RT. After three washes with 0.1M cacodylate buffer, the samples were post-fixed with 1% OsO4 for 1h at RT. Then the samples were dehydrated in sequential ethanol solutions and embedded in an Eponate mixture (Electron Microscopy Sciences, Hatfield, PA, USA) for polymerization for 24h at 60°C. Ultrathin sections (∼70nm) were cut with a diamond knife. The sections were re-stained with uranyl acetate and lead citrate and then photographed using a transmission electron microscope (FEI Tecnai G2 12, Holland). For the calculation of the ERMICC contact index, the formula is the LIN/(PerM×DistER-M). LIN represents the interface length between mitochondria and ER; PerM indicates mitochondria perimeter, and DistER-M is the distance between mitochondria and ER. About 100 mitochondria were considered to calculate for Mito-ER distance, Mito-ER contacts, and ERMICC contact index.

### Cell culture and transfection

HEK293T (293T) cells were cultured in DMEM medium with 10% serum. Transfection performed using Lipofectamine 2000 (Invitrogen) according to the manufacturer’s instructions. For the transfection of plasmid, 4μ used in a 45mm diameter plate.

### Sertoli cell purification

For Sertoli cell isolation, testes from adult mice were dissected in DHANKS medium and incubated in DMEM/F12 medium containing 1 mg/ml collagenase IV, 0.5 mg/ml Deoxyribonuclease I and 0.5 mg/mL hyaluronidase for 10min at 37 °C. After centrifuge (1000 rpm/min), the precipitates were collected and incubated in the DMEM/F12 medium with 2.5 mg/ml trypsin and 0.5 mg/ml Deoxyribonuclease I for 10min at 37 °C. Then seminiferous tubules were incubated with DMEM/F12 containing 10%FBS for 5 min to inhibit the trypsin digestion. Subsequently, Sertoli cells collected after filtering through 40 μm pore-size nylon mesh and stored for the experiments.

### Protein extracts and Western blots

Protein extracts were prepared from mouse tissues using RIPA lysis buffer (150 mM sodium chloride, 1.0% NP-40, 0.5% sodium deoxycholate, 0.1% SDS, and 50mM Tris, pH=8.0). Protein extracts were denatured with 2X loading buffer (4% SDS, 10% 2-mercaptoethanol, 20% glycerol, 0.004% bromophenol blue, 0.125M Tris-HCl, pH=6.8) at 95°C for 10 min and run on a 10% SDS-PAGE, followed by transfer to PVDF membrane. The membrane was probed with primary antibodies (Supplementary Table S7) followed by secondary antibody treatment at 1:10000 dilutions (anti-mouse HRP and anti-rabbit HRP, abbkine). Immun-Star™ HRP (1705040, BIO-RAD) was used for chemiluminescence detection and photographed by ChemiDoc XRS+ system (BIO-RAD).

### Co-immunoprecipitation

For *in vivo* co-immunoprecipitation, mouse testes were homogenized in lysis buffer (20mM HEPES, pH=7.3, 150mM NaCl, 2.5mM MgCl_2_, 0.2 % NP-40, and 1mM DTT) with protease inhibitor cocktail (P1010, Beyotime, China). Relevant antibody and pre-cleaned magnetic protein A/G beads added to the tissue lysate. The lysate was incubated overnight at 4°C with gentle agitation to form the immunocomplex, and the magnetic beads were washed using the lysis buffer for 5 times. The pellet was re-suspended with 30μl lysis buffer and 30μl 2X loading buffer (4% SDS, 10% 2-mercaptoethanol, 20% glycerol, 0.004% bromophenol blue, 0.125 M Tris-HCl, pH=6.8), pipetted up and down several times to mix and elute the protein from the beads, then boiled at 95°C for 10 min and run on a 10% SDS-PAGE. For in vitro co-immunoprecipitation, full-length *Mfn2* cDNA was cloned into a modified pcDNA3 vector encoding a 2X FLAG-tag, and full-length *Miwi, Ddx4*, *Msy2*, and *Tdrkh* cDNAs were cloned into pCMV vector carrying an MYC-tag. HEK293T cells were transfected with indicated plasmids using Lipofectamine 2000 (Life Technologies). After 48h, immunoprecipitation was performed using anti-MYC rabbit polyclonal antibody (10828-1-AP, Proteintech), followed by a Western blot to identify protein interactions.

### RNA isolation, reverse transcription and RT-qPCR

Total RNA was extracted from mouse tissues using Trizol reagent (Invitrogen, Cat no.15596-025) following the manufacturer’s instructions. For complementary DNA (cDNA) synthesis, 1μ g of RNA was treated with DNase I (Progema, M6101) to remove the residual genomic DNA and reverse transcribed with cDNA Synthesis Kit (ThermoFisher, Cat no. 4368814). RT-qPCR was performed using an SYBR Premix on a StepOne Plus machine (ABI). Relative gene expression was analyzed using the 2^-Ct^ method with *Arbp* as an internal control. All primers are shown in Supplementary Table S8.

### Purification of spermatogenic cells

STA-PUT method based on sedimentation velocity at unit gravity (Bellve et al., 1977; Bellve, 1993) was used to purify the pachytene spermatocytes and round spermatids from adult WT and *Mfn2*-cKO mouse testes. We assessed cellular morphology using phase contrast microscopy to determine the purity of the cell population. Pachytene spermatocytes and round spermatids with ≥90% purity were used for RNA-seq analyses.

### RNA sequencing and bioinformatics analysis

After purification of pachytene spermatocytes and round spermatids from WT and *Mfn2*-cKO testes, total RNA was isolated using the Trizol reagent following the manufacturer’s protocol and treated with DNase I to digest residual genomic RNA. The purity, concentration, and integrity were assessed using a NanoDrop 2000 spectrophotometer (Thermo Scientific), a Qubit RNA Assay Kit in Qubit 2.0 Fluorometer (Life Technologies), and a Nano 6,000 Assay Kit of the Bioanalyzer 2,100 system (Agilent Technologies), respectively. Then, a total amount of 2µg of RNA per sample was used to prepare poly(A+)-enriched cDNA libraries using the NEBNext Ultra RNA library Prep Kit for Illumina (New England BioLabs) according to the manufacturer’s instructions, and base pairs (raw data) were generated by the Illumina Hi-Seq 2500 platform. Raw reads were processed with cutadapt v1.9.1 to remove adaptors and perform quality trimming. Trimmed reads were mapped to the UCSC mm10 assembly using HiSAT2 (V2.0.1) with default parameters. Differentially expressed genes for all pairwise comparisons were assessed by DESeq2 (v1.10.1) with internal normalization of reads to account for library size and RNA composition bias. Differentially regulated genes in the DESeq2 analysis were defined as two-fold changes with an adjusted *Ρ*-value of <0.05. Alternative splicing (AS) patterns were processed with rMATs software (replicate Multivariate Analysis of Transcript Splicing)(Shen et al., 2014). Splicing events with FDR (false discovery rate) <0.05 and |Δψ|>0.05 were defined as misregulated AS events. Gene Ontology (GO) and Kyoto Encyclopedia of Genes and Genomes (KEGG) analyses were conducted using the database for annotation, visualization, and integrated discovery (DAVID)(Dennis et al., 2003). Rich factor=(number of differentially expressed genes in GO term)/(total number of genes in GO term). The larger rich factor is, the higher enrichment is. The integrative genomics browser tool (IGB)(Freese et al., 2016) was used for efficient and flexible visualization and exploration of spliced sites between WT and *Mfn2*-cKO samples on standard desktop computers.

### Small RNA-Seq library preparation and sequencing

For preparing small RNA-seq libraries, two biological testis samples at P25 from WT and *Mfn2*-cKO mice were collected, respectively. Total RNAs extracted using TRIzol reagent (Invitrogen) and subsequently processed to prepare small RNA-Seq library and sequencing as previously described(Dong et al., 2019). For the length distribution of piRNAs analysis, data were normalized by miRNA reads (21–23nt) from small RNA sequencing.

### Sucrose density gradient fractionation and polysome profiling

Testicular seminiferous tubules were dissected and collected according to previous literature with minor modifications(Aboulhouda et al., 2017; Karamysheva et al., 2018). Briefly after adding polysome extraction buffer (20mM Tris-HCl, pH 7.4, 100 mM KCl, 5 mM MgCl_2_, 1 mM DTT, 0.5% NP-40, 1x protease inhibitor cocktail (EDTA-free), 200 μg/ml CHX, 200 units/mL of RNase inhibitor), the tubules were transferred to a small (0.5-1.0 mL) Dounce homogenizer to disrupt the tissues with seven to eight strokes of the glass pestle. The samples were centrifuged at 12,000g in 4°C for 10 min to collect the supernatant into a new tube. Then, DNase I (final 5 U/ml) was added into the tube, and the tube was placed on ice for 30 min to allow the DNase I to degrade any DNA contamination. Equal amounts of cytoplasmic lysates were carefully loaded on the top of a 10-50% sucrose gradient. All tubes were equally balanced, and then the gradients were centrifuged for 3h in SW41Ti swinging bucket rotor at 160,000g at 4 °C using a Beckman ultracentrifuge. Fractions (300 μl/tube) were collected manually and immediately transferred to an ice bucket. The absorbance was detected at 260nm to display the polysome profile of the gradients. The proteins were precipitated with 9 volumes of 96% ethanol.

### Statistical analysis

All data are presented as mean ± SEM unless otherwise noted in the figure legends. Statistical differences between datasets were assessed by one-way ANOVA or Student’s t-test using the SPSS16.0 software. *P*-values are denoted in figures by *, *P* < 0.05; **, *P* < 0.01; and ***, *P* < 0.001.

### Data availability

All data needed to evaluate the conclusions in the paper are present in the paper and/or the Supplementary Materials. All RNA sequencing data are deposited in the NCBI SRA (Sequence Read Achieve) database with the accession number of SRP212036. All other supporting data of this study are available from the corresponding author upon reasonable request.

## Acknowledgments

We are grateful for engaging discussions with colleagues from Huazhong University Science and Technology, China, in the very initial phase of the project. This work supported by grants from National Natural Science Foundation of China (31671551 and 81971444 to S.Y.), the Science Technology and Innovation Commission of Shenzhen Municipality (JCYJ20170244 to S.Y.), Natural Science Foundation of Hubei Province (2017CFA069 to S.Y.).

## Author contributions

X.W. and S.Y. conceived and designed the research. X.W., Y.W., J.Z., S.G., C.C., S.K., and Z.Z. performed all bench experiments and data analyses. S.Y., X.W., and Z.Z. wrote the manuscript. S.Y. supervised the project. All authors read and approved the manuscript.

## Conflict of interest

The authors declare that they have no conflict of interest.

